# Pathway-Level Information ExtractoR (PLIER) for gene expression data

**DOI:** 10.1101/116061

**Authors:** Weiguang Mao, Elena Zaslavsky, Boris M. Hartmann, Stuart C. Sealfon, Maria Chikina

## Abstract

A major challenge in gene expression analysis is to accurately infer relevant biological insight, such as regulation of cell type proportion or pathways, from global gene expression studies. We present a general solution for this problem that outperforms available cell proportion inference algorithms, and is more widely useful to automatically identify specific pathways that regulate gene expression. Our method improves replicability and biological insight when applied to trans-eQTL identification.

## 1 Introduction

One salient feature of high dimensional molecular data structure is the presence of groups of correlated measurements. In gene expression datasets, correlation among genes commonly represents coordinated transcriptional regulation or, in studies of heterogeneous tissues, variation in cell-type proportion. Identifying the mechanisms underlying coordinated regulatory changes, such as the activity of upstream regulatory pathways, is crucial for interpretation. Furthermore, determining changes in tissue composition across samples is important for identifying experimental effects on cell type proportion and for improving statistical power for identifying gene regulation occurring within cell subtypes. Notably, correlated expression patterns may also be the result of various technical factors, often referred to as “batch effects” (see Leek et al. [2010] for review). The challenge is to identify and interpret biologically meaningful signatures while reducing any negative effects of technical noise. To meet these goals, we have developed Pathway-Level Information ExtractoR (PLIER). PLIER performs an unsupervised data structure deconvolution and mapping to external knowledge, reducing noise and identifying regulation in cell-type proportions or pathways.

PLIER approximates the expression pattern of every gene as a linear combination of eigengene-like latent variables (LVs). In constructing LVs, PLIER surveys a large compendium of prior knowledge (genesets) and produces a dataset deconvolution that optimizes alignment of LVs to a relevant subset of the available genesets. The method automatically finds these relevant genesets among the hundreds to thousands considered (see Fig. 1A). Technical noise reduction is also achieved during the deconvolution by capturing these batch-effect correlated gene changes in some of the LVs that are not associated with genesets. By integrating dataset deconvolution with prior knowledge, this process reduces technical noise, increases statistical power, and identifies the specific upstream pathways or cell type proportion changes driving the geneset-aligned signals.

**Figure 1:**
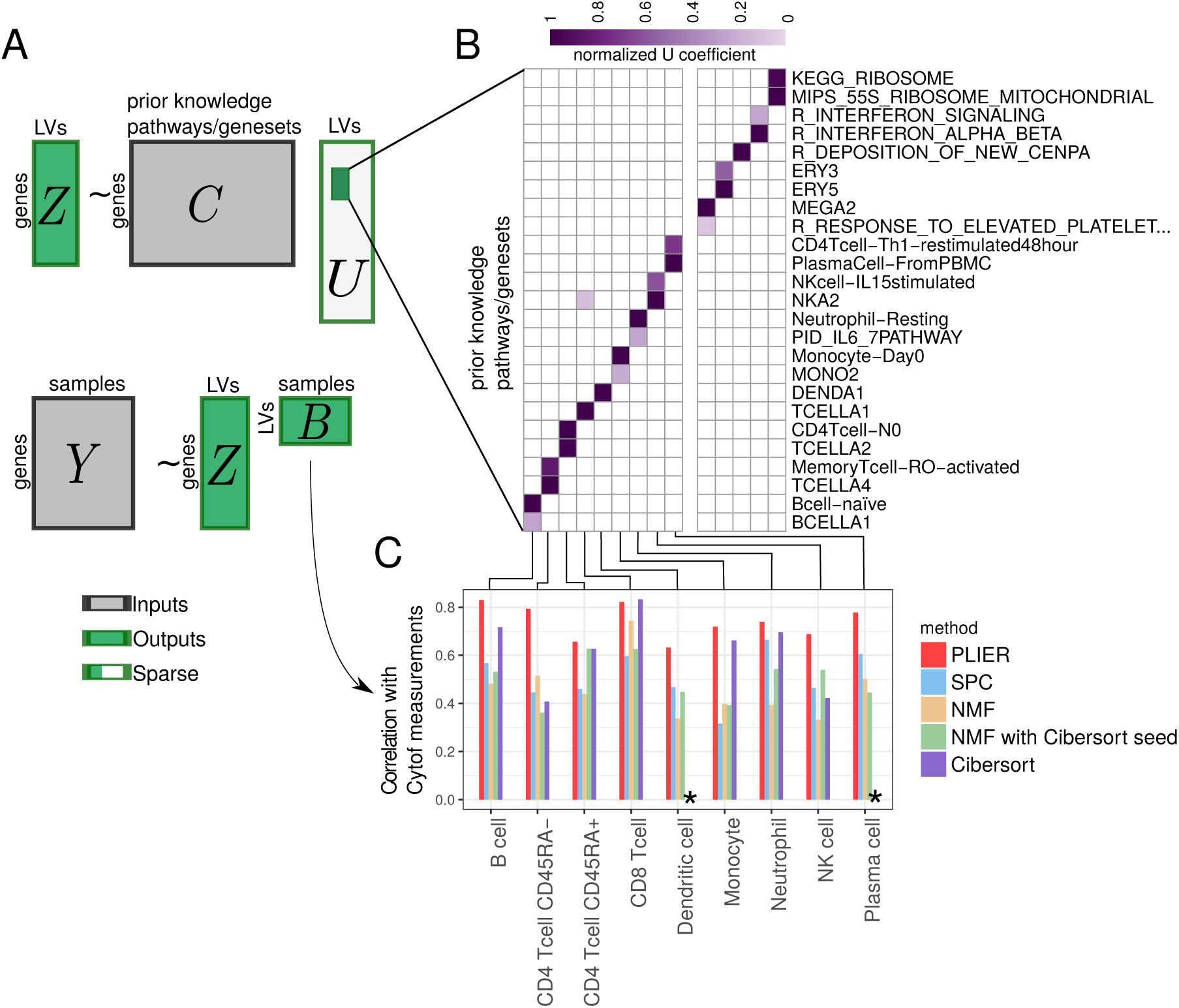
PLIER overview. PLIER is a matrix factorization approach that decomposes gene expression data into a product of a small number of latent variables and their corresponding gene associations or loadings, while constraining the loadings to align with the most relevant automatically selected subset of prior knowledge. Given two inputs, the gene expression matrix *Y* and the prior knowledge (represented as binary geneset membership in matrix C), the method returns the latent variables (*B*), their loadings (*Z*), and an additional sparse matrix (*U*) that specifies which (if any) prior information genesets and pathways are used for each latent variable. The light gray area of *U* indicates the large number of zero elements of the matrix. We apply our method to a whole blood human gene expression dataset. The positive entries of the resulting *U* matrix are visualized as a heatmap, facilitating the identification of the correspondence between specific latent variables deconvolved from the data and prior biological knowledge-based geneset categories. We validate the latent variables mapped to specific cell-types by comparing PLIER estimated relative cell-type proportions with direct measurements by Mass Cytometry. We find that the PLIER estimates are highly accurate, outperforming other matrix decomposition methods as well as the state-of-the-art dedicated blood mixture deconvolution method, Cibersort [Newman et al., 2015]. (*) indicates that the Cibersort estimate had near zero negative correlation with the directly measured proportion for these cell-types.

## 2 Results

### 2.1 Estimating Blood Composition Variables

We first validate the method using cell-type proportion inference because it is an important objective, other methods are available for comparative benchmarking, and predictions can be tested against a direct measurement gold standard. For this purpose, we generated a validation dataset comprising 35 human whole-blood samples assayed both by RNA-seq and direct CYTOF measurement of cell type proportion. We applied PLIER to the validation dataset using 605 pathways which included 60 cell-type markers and 555 canonical pathways from MSigDB. We produced a decomposition with 14 latent variables annotated with high confidence (AUC >0.7, FDR < 0.05, see methods for cross-validation procedure) to one or more genesets, of which 9 represented cell types also measured by the Cytof panel. The correlation between the cell type PLIER LVs and Cytof measurements in these 35 samples had a mean of 0.76 (range 0.63-0.84) (Fig. 1).

We compared PLIER against the established current methods for mixture decomposition inference. These methods either rely on low-rank matrix decomposition or model-based methods that fit gene expression values to reference signatures. We include the most widely used constrained matrix decomposition approaches: non-negative matrix factorization (NMF), and Sparse Principle Component Analysis (SPC) (see Methods for details). For a model-based approach, we tested Cibersort [Newman et al., 2015] because it has been shown to be superior to other model-based methods. We also tested a hybrid decomposition/model-based approach which is implemented by seeding the NMF decomposition with known pure cell expression values taken from the Cibersort reference signature for human blood.

PLIER outperformed all constrained matrix decompositions, including the seeded NMF. Surprisingly, for 7 out of the 9 celltypes, PLIER also outperformed Cibersort, a supervised method explicitly developed for this problem. The excellent performance of the essentially unsupervised and general PLIER method in comparison with Cibersort most likely results from the capacity of PLIER to sort through many candidate genesets and find the ones most informative for the specific data structure of the dataset under study. PLIER can be supplied with multiple and even discordant markers sets for the same cell-type. Indeed, we find that while both the IRIS and DMAP prior knowledge datasets included in this analysis contain neutrophil and dendritic cell genesets, only the markers derived from IRIS were selected to represent neutrophils in this deconvolution, while the markers for dendritic cells came from DMAP. The biological validity of this varied geneset selection is supported by the excellent correlation with direct cell type proportion measurements.

### 2.2 Application to trans-eQTL discovery

While PLIER shows excellent performance when benchmarked for cell type deconvolution, it is applicable for a wide variety of problems in integrative biological interpretation of gene expression data, including pathway analysis. By aligning gene expression patterns in a dataset under analysis with prior information genesets, PLIER produces a decomposition that isolates biological pathway level effects as well as technical and systemic variation into separate components. By simultaneously optimizing and isolating latent variables matched to genesets and to technical artifacts, PLIER reduces noise and increases power to detect true and distinct pathway-level effects. Another important feature of PLIER is that it automates these interpretive processes in an easy to use, accessible framework.

As an example, we evaluated the usefulness of PLIER for the difficult task of genotype-quantitative trait association. Two groups of eQTLs are typically distinguished: locally acting *cis*-eQTLs that affect a nearby gene, and *trans*-eQTLs that are commonly mediated at the pathway level [Battle et al., 2014]. Many *trans*-eQTLs exert their effect by altering the activity of a regulatory protein, which in turn affects the expression of many downstream genes [reference Lude et al.]. *Trans*-eQTLs, which provide important insight into gene regulatory networks, are difficult to detect and are less commonly identified than *cis*-eQTLS.

We analyze the recently published DGN dataset [Battle et al., 2014], which contains whole blood RNAseq and genotype measurements from 922 individuals, to demonstrate how the PLIER framework extracts a broad spectrum of pathway effects and enables network-level eQTL discovery and interpretation. For the candidate prior information, we used a comprehensive collection of 4445 genesets comprising biochemical and transcriptional pathways (“canonical pathways” and “chemical and genetic perturbations” from MSigDB [Subramanian et al., 2005]), cell-type markers from multiple sources [Newman et al., 2015, Abbas et al., 2009, Novershtern et al., 2011] and cytokine signatures [Filiano et al., 2016]. The PLIER decomposition produced 86 LVs that have at least one matched pathway with an FDR > 0.05, and were associated overall with 318 of the 4444 pathway genesets evaluated. The decomposition captured cell-type variation with a high degree of specificity, differentiating naive and memory B-cells, plasmacytoid and myeloid dendritic cells, regulatory T cells and multiple subtypes of CD8 T-cells. PLIER also captured variation in non leukocyte celltypes such as megakaryocytes and erythrocytes, and transcriptionally mediated pathway effects such as Type I and Type II interferon signaling, and NKFB pathway. Overall, based on geneset utilization and top associated genes, 29 LVs were unambiguously related to cell-type, canonical pathways or cytokine signaling (see Figure 8 for *U* matrix visualization and Supplementary File 1 for a complete list of LV-geneset associations).

**Figure 8:**
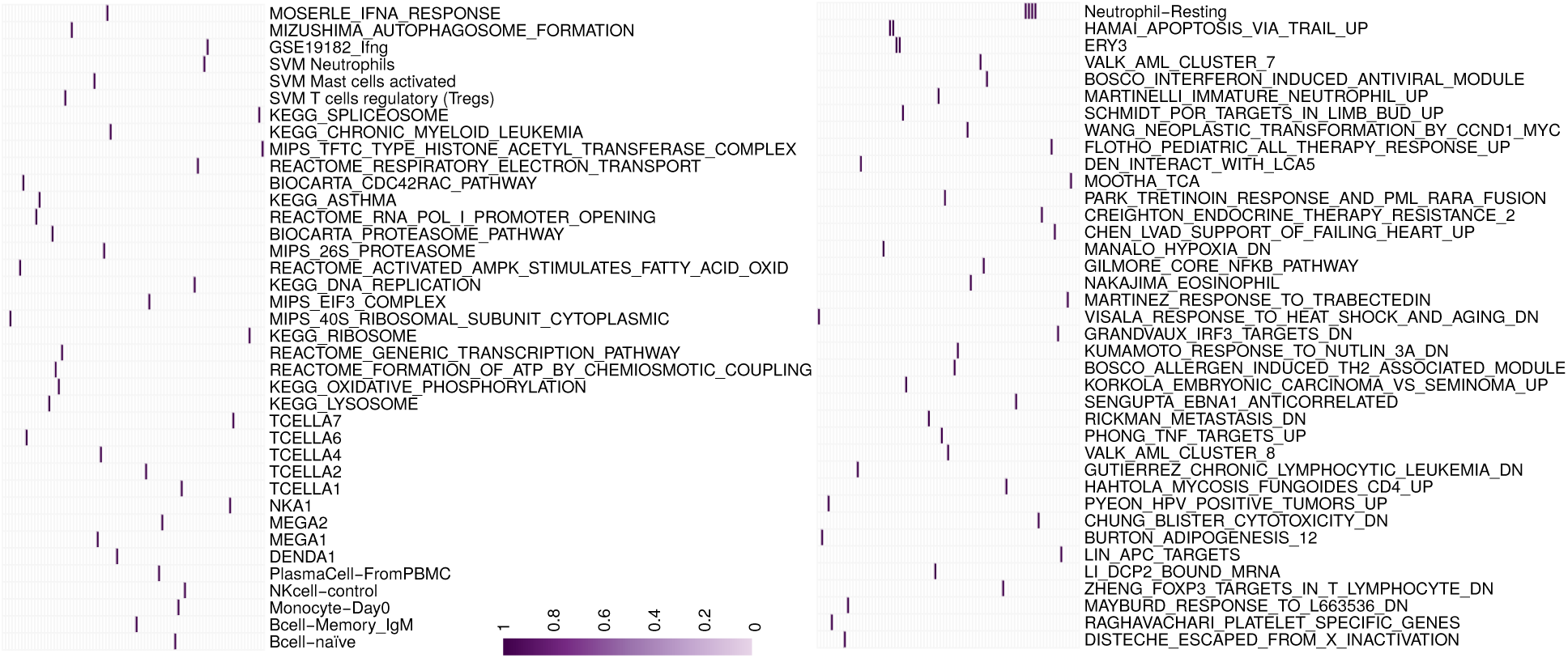
The *U* matrix for the complete set of LVs that had at least one pathway match with and FDR<0.05. For visualization, only the top pathway (largest *U* coefficient) is shown. We note that pathways for abundant cell-types such as neutrophils and erythrocytes are top hits for multiple LVs.

In order to perform eQTL analysis, we treated the PLIER LVs as quantitative traits, and identified 14 LVs showing significant associations with genotypes (Table 1). In contrast to gene level QTL identification, the PLIER LV QTLs are pathway level eQTLs that capture the concerted behavior of multiple genes (Fig. 2A). The gold-standard for eQTL discovery is reproducibility in an independent dataset. We compared the replication rates of standard gene-level trans-eQTLs and PLIER-identified pathway eQTLs using an independent paired blood RNA-seq GWAS dataset (NESDA, (REF)). We compared the replication rates of PLIER identified pathway-level eQTLs and gene-level eQTLs by determining the true positive rate π_1_ (see Methods). When we examine all gene-level trans-eQTLs in the NESDA dataset, we find that π_1_ is 0.36. In contrast, the same analysis on the pathway eQTLs identified by PLIER showed a higher π_1_ value of 0.6. Using an FDR threshold of 0.05 we found that 0.58 (51 out of 88) of PLIER eQTLs were replicated in contrast to gene-level replication of 0.22 (67 out of 309). These results suggest that the improved replicability of PLIER pathway-level eQTLs results from the QTL assignment being based on an integrated, biologically relevant signal captured in the pathway LV eQTL.

**Table 1:** Summary table of all pathway-level effects found in the DGN dataset. Indicated p-values are Bonferroni corrected for the total number of SNPs (651075). In most cases pathways were named based on their geneset association captured in the *U* matrix. Some pathways are named based on further analysis of the expression patterns of top gene in a independent dataset of mouse immune cells, ImmGen [Heng et al., 2008] (See Figure 10) and/or a the presence of a putative *cis* eQTL transcriptional mediator. The complete pathway utilization for these LVs can be seen in Figure 2. The expression patterns for top 15 genes driving each latent variable are plotted in Figure 9. Latent variables with no pathway association in PLIER decomposition (that is no positive entries in *U*) are starred.

| LV id | LV name | snps | cis-Gene(s) | corrected p-value |
| --- | --- | --- | --- | --- |
| 44 | Mega/platelet 1 | rs1354034 | ARHGEF3 | < 1.45e-10 |
| 133 | Mega/platelet 2 | rs1354034 | ARHGEF3 | 0.01547 |
| 120 | Histones | rs1354034 | ARHGEF3 | 0.01889 |
| 97 | Zinc fingers, pseudogenes | rs1471738 | SENP7 | < 1.45e-10 |
| 56 | PLAGL1 associated, myeloid | rs9321957 | PLAGL1 | 3.6e-05 |
| 42* | IKZF1 associated, myeloid | rs10251980 | IKZF1 | < 1.45e-10 |
| 17 | NEK6 associated, myeloid | rs16927294 | NEK6 | 0.00360 |
| 67 | Neutrophils | rs13289095 | PKN3,SET,ZDHHC12 | 0.01888 |
| 55* | NFE2 associated, erythrocyte | rs35979828 | NFE2 | < 1.45e-10 |
| 21 | Interferon-gamma | rs3184504 | SH2B3 | 5.9e-05 |
| 40 | NFKB/TNF | rs12100841 | PPP2R3C | 0.00204 |
| 16 | Myeloid/ILC | rs1138358 | BCL2A1,MTHFS,ST20 | 0.00025 |

**Figure 2:**
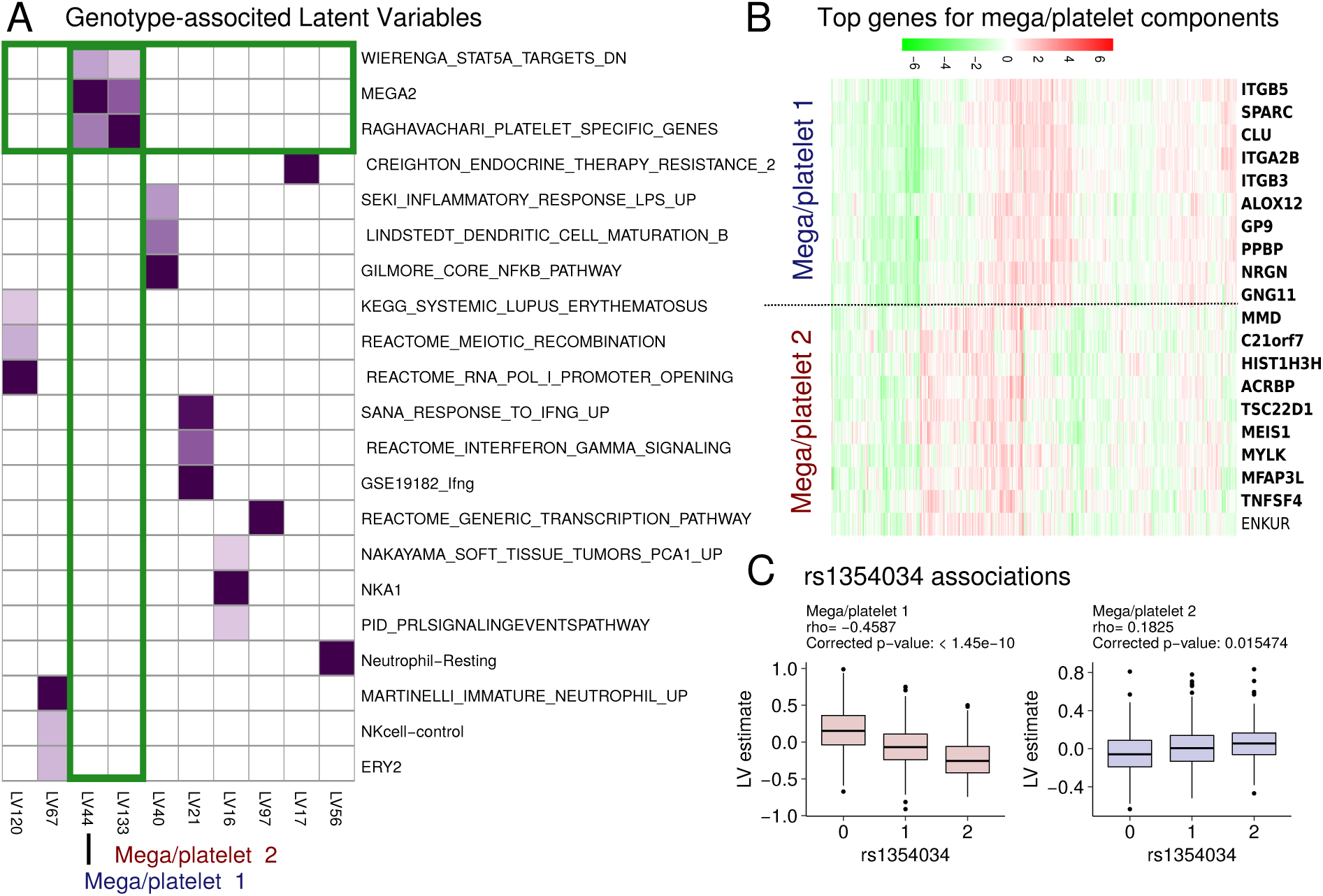
**A** A heatmap of a subset of the *U* matrix corresponding to LVs with a genotype effect (LV eQTLs). Only pathways with a cross-validation FDR of < 0.05 are shown. We find that two latent variables (LV44 and LV133) share pathway annotations (albeit with different coefficient) that suggest a relationship with megakaryocyte and platelet biology. **B** Heatmap of the top genes in the loading for LV44 and LV133. Genes that are annotated to the pathways shown in panel A are in bold. **C** Boxplots of the association of LV44 and LV133 with SNP rs1354034. While the LV estimates are positively correlated, the effects of rs1354034 are opposite. These results indicate that the pathways captured by the expression patterns of LV44 and LV133 are independently regulated by the rs1354034 locus.

Besides improving the accuracy of eQTL discovery, the PLIER decomposition identifies the pathway(s) associated with the LV eQTL, which can provide precise biological interpretation of the genetically regulated processes. For example, PLIER shows that SNP rs1354034 (located within gene ARHGEF3) is associated with two LVs, LV44 and LV133, that are related to megakaryocyte/platelet lineage based on their pathway association (Fig. 2A,B). In the published gene level analysis of the DGN dataset, this SNP yields the largest number of significant *trans*-eQTLs, however no biological interpretation was inferred [Battle et al., 2014]. Using PLIER, we find that both associated LVs are annotated to platelet pathway processes, which is consistent with a known effect of this SNP on platelet number (PLT) and platelet volume (MPV) [Gieger et al., 2011]. However, our analysis further shows that the two LVs linked to this SNP are supported by different genes that show distinct expression patterns (Fig. 2B). These results suggest that the two LV eQTLS may distinguish two biologically distinct aspects of platelet/megakaryocyte biology. A recent hematopoetic lineage report supports this formulation. This single cell study shows that genes associated with the two LVs express at different developmental time points [Olsson et al., 2016]. Specifically, mouse orthologs of MEIS1 and TSC22D1 (from LV133) are expressed in all megakaryocyte precursors, while ITGA2B (from LV44) is megakaryocyte specific, suggesting that these two LVs capture processes that are active at different times in megakaryocyte development.

LV133 and LV44 are positively correlated with each other in the DGN dataset. Notably, the effects of the rs1354034 alleles on LV133 and on LV44 go in opposite directions (Figure 2C). Furthermore, we find that using partial correlation analysis, whereby the LVs are corrected for each other, dramatically improved the eQTL statistics (Figure 11). These results strongly argue that the LV44 and LV133 effects are independent.

**Figure 11:**
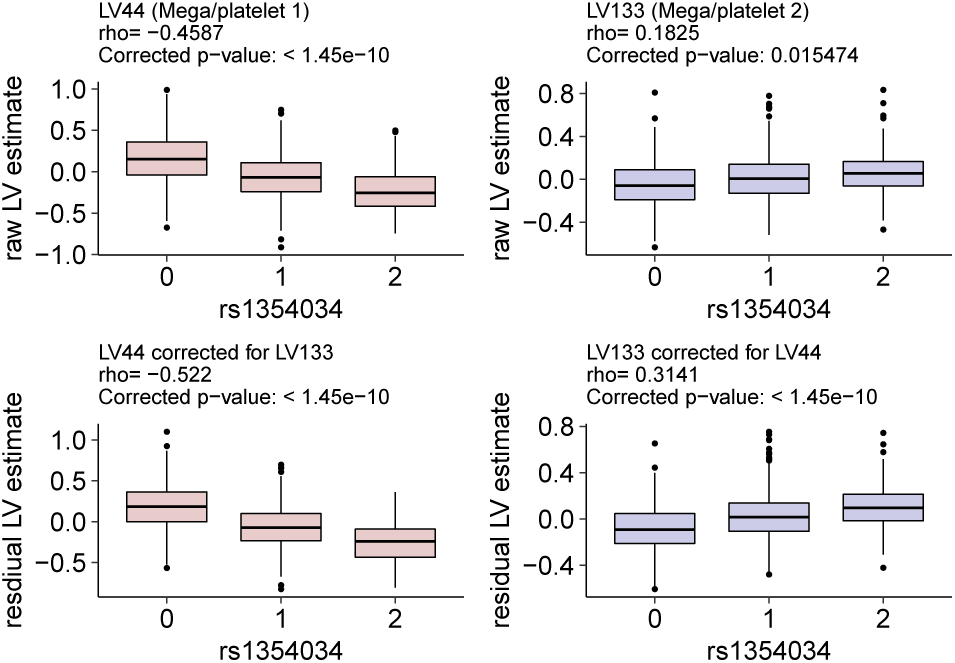
The effects of rs1354034 of LVs 44 and 133. First row, using raw estimates. Second row, using corrected estimates (residuals from the linear regression fit).

We speculate that the independent regulation of the two LV eQTLs by the same locus results from an effect on different specific regulator effectors (e.g. transcription factors) that are modulated by rs135034 at different periods of megakaryocyte development. The rs1354034 SNP is known to be pleiotropic as it is linked to both MPV and PLT phenotypes, which are affected independently by other genetic variation [Gieger et al., 2011]. We hypothesize that the effects of rs1354034 on multiple LVs is reflective of its pleiotropic function. Indeed, analysis of the association of the two LVs with SNPs known to be specifically linked to MPV or PLT alone shows divergent patterns. We find that in addition to the association with rs1354034, the developmentally early LV133 is most strongly associated with a SNP linked to platelet number, whereas the later LV44 is most strongly associated with a SNP linked to platelet volume (Table 2). This analysis supports a model where ARGHEF3 exerts its pleiotropic affects on platelet volume and number at different developmental time points. These results demonstrate how PLIER can leverage dataset structure and external knowledge to resolve fine-grained mechanistic insight underlying complex biological processes.

**Table 2:** Summary table of the associations between the two mega/platelet LVs and SNPs known to affect only one platelet phenotype. A total of 80 SNPs with known platelet phenotypes were tested [Gieger et al., 2011]. While no SNPs outside of the ARGHEF3 locus achieved genome-wide significance, some associations were significant when applying a Bonferroni correction only to the 80 platelet specific hypotheses (significant p-values are in bold). We find that the associations of the two mega/platelet LVs with other loci known to affect platelet biology are distinct. Our analysis suggests that the early mega/platelet LV (LV133) is more closely related to the process controlling platelet number (PLT) while the late mega/platelet LV (LV44) is related to the process controlling platelet volume (MPV).

| phenotype | reported SNP | Close gene | LV44 p-value | LV 133 p-value | proxy SNP |
| --- | --- | --- | --- | --- | --- |
| MPV | rs10876550 | COPZ1 | **1.1847e-05** | 0.69933 | rs10876550 |
| PLT | rs2911132 | ERAP2 | 0.13817361 | **2.4417e-05** | rs2549803 |

### 2.3 Simulation

Since in a real dataset the true data generating model is unknown and is likely more complex than what can be captured with a dimensionality reducing matrix decomposition, we use a simulation to evaluate the operating characteristics of our method. We hypothesize that our method is able to more accurately recover the “correct” LVs by rotating the matrix decomposition to align with prior knowledge.

We simulate data with 5000 genes, 300 samples, and 30 latent variable according to the NMF model.

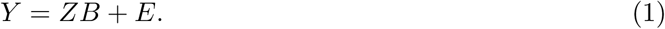

With both *Z* and *B* > 0. *B* is drawn from Beta distribution and each column sums to one by design. The columns of *Z* are drawn from Gamma distribution Γ(5,1). The matrix *E* ∈ 𝒩(0,1) represents random noise. We also generate a prior knowledge matrix *C*. For each column of *Z*, we randomly pick up a threshold value on the percentage of genes which belong to a hypothetical prior knowledge geneset. The threshold value varies from 0.01 to 0.1 with a step size 0.01, which is in consistent with that of real biological genesets. With the threshold value, we select the corresponding fraction of genes which come with top values in the column of *Z* to construct the prior knowledge geneset. Also we generate additional uninformative genesets by randomly picking genes. For the purpose of applying PLIER and SPC the final data is z-scored.

Our basic evaluation strategy is based on computing the maximal correlations between simulated and recovered latent variables, for the purpose of comparisons with other methods we use the absolute value so as to allow factors with reversed sign. Figure 3 depicts the results of multiple simulation runs process with four decomposition methods: PLIER, PLIER with no prior information (which can be accomplished by setting λ_3_ to a high value), NMF[Brunet et al., 2004] and SPC [Witten et al., 2009]. NMF is a popular decomposition method that is free of hyperparameters (though different matrix norms can be used) however it requires positive data as input. SPC is another popular method that can enforce sparsity and positivity, it has one hyperparameter that we set by cross-validation for each component as described in the original paper [Witten et al., 2009]. Among these methods only PLIER is able to reliably produce high correlations with the simulated latent variables and only when using prior information. Importantly, we emphasize that the simulation is not based on a PLIER model where we assume that loadings of genes in the pathway and outside the pathway differ by a constant factor but is rather based on the NMF model. Nevertheless the PLIER approach is effective even in the case where the model design differs from the underlying assumptions.

**Figure 3:**
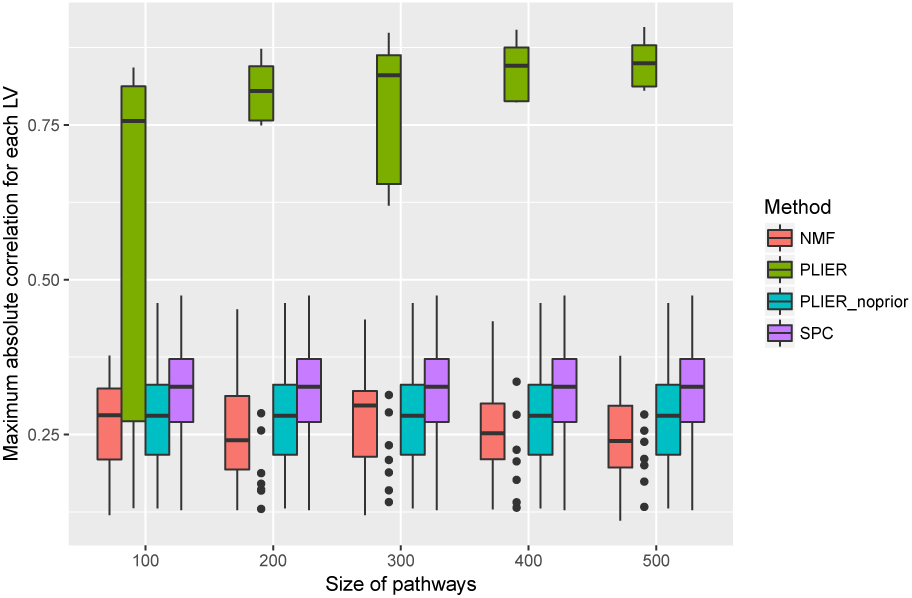
Data is simulated according to the NMF model (see text for details). Boxplots of the correlation between simulated LVs and those recovered by various decomposition methods. We compare PLIER against two other methods, NMF [Brunet et al., 2004] and SPC [Witten et al., 2009], as well as PLIER run without using any prior information. In this simulation we provide PLIER with 1000 pathways of which only 30 are correct and vary the size of the prior information pathways provided to PLIER. We find that the best performance is achieved by PLIER specifically when prior information is used with a notable improvement when prior information pathways are larger.

We also investigate how adding noise to the prior information affects performance, hypothesizing that as more irrelevant geneset are included in our prior knowledge matrix *C*, the advantage of using prior information will be reduced. Repeating the experiment above with varying sets of noninformative pathways we find that the performance indeed drops off as the total number of pathways is increased to 10,000 though even at that level of prior information noise PLIER outperforms other methods (Figure 4).

**Figure 4:**
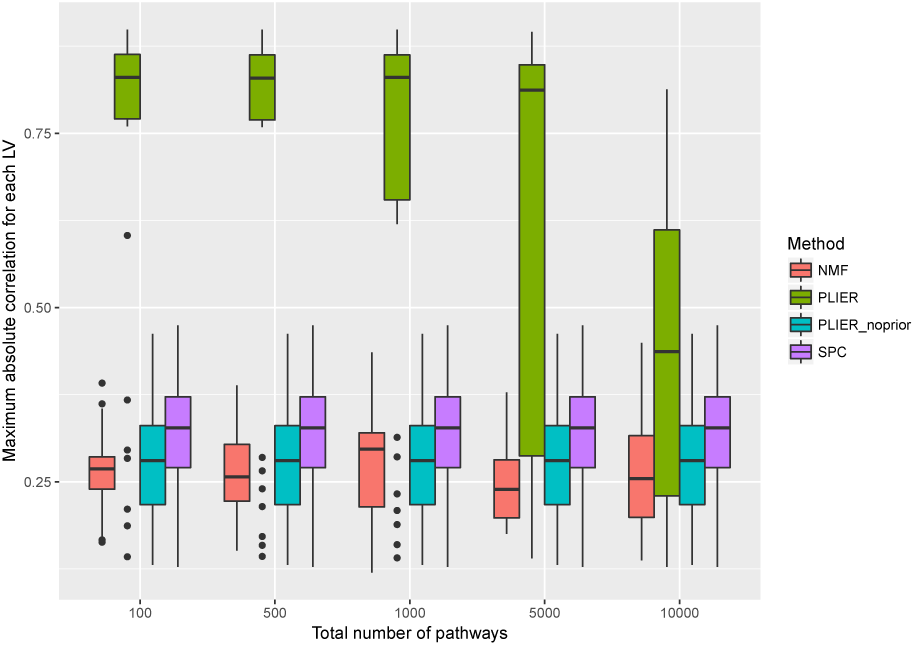
Data is simulated as in Figure 3 except that the number of genes per pathway is kept at 300 and the number of uninformative pathways is varied. As the prior information gets noisy PLIER performance approaches that of other We also investigate how adding noise to the prior information affects performance, hypothesizing that as more irrelevant geneset are included in our prior knowledge matrix *C*, the advantage of using prior information will be reduced. Repeating the experiment with varying sets of noninformative pathways we find that the performance indeed drops off as the total number of pathways is increased to 10,000 though even at that level of prior information noise PLIER outperforms other methods (Figure 4).constrained decomposition methods.

**Figure 5:**
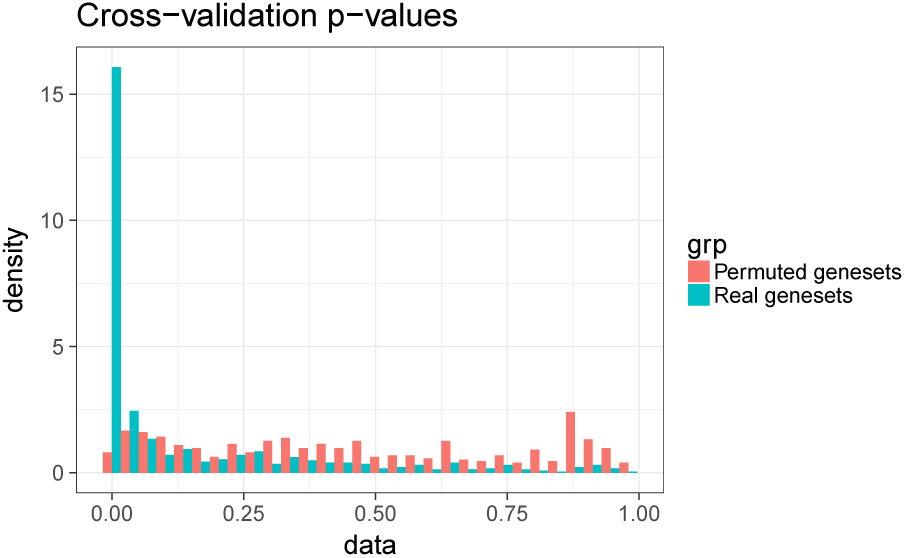
Histogram comparisons of pathway association p-values produced with real genesets and a single run with gene-label permuted genesets.

### 2.4 Pathway recovery significance

We estimate the significance of LV pathway association by removing a random 1/5 of the genes annotated to each pathway prior to running PLIER. For each LV-pathway correspondence represented as a positive value in *U* we compute the AUC and p-value for the recovery of that pathway in the loadings of *Z* using the held-out set of genes as positive labels and genes not annotated to this pathway as negative labels. We verify that this procedure produces correct estimates by running PLIER with a the geneset collection used for the DGN dataset but randomly permuted gene labels. Gene-level permutation preserves the pathway size distribution and dependency structure but should not have any non-random associations with the structure of the gene expression dataset. We find that in the permuted setting our cross-validation procedure produces uniformly distributed p-values.

### 2.5 Technical variation invariance

A key motivation for PLIER is to tease apart technical and biological variation. Specifically, the hypothesis is that LVs that use prior information are indeed of biological origin. If that is the case we expect that PLIER results are relatively insensitive to normalization for technical factors and we test this hypothesis by applying PLIER to differently normalized data. The DGN datasets [Battle et al., 2014] used in this study has been normalized for 20 technical variables which reflected information about data collection and RNAseq quality control. We can also apply PLIER to the “naive-normalized” version of the same data represented by log-transformed counts normalized by quantile normalization. Obtaining two different decompositions we find that many LVs can be matched in one-to-one correspondence based on rank-correlation of the loadings. Correlations for top matched pairs are show in Figure 6. Moreover, the matching LVs use prior information genesets that are either the same or closely related (see row/column names in Figure 6).

**Figure 6:**
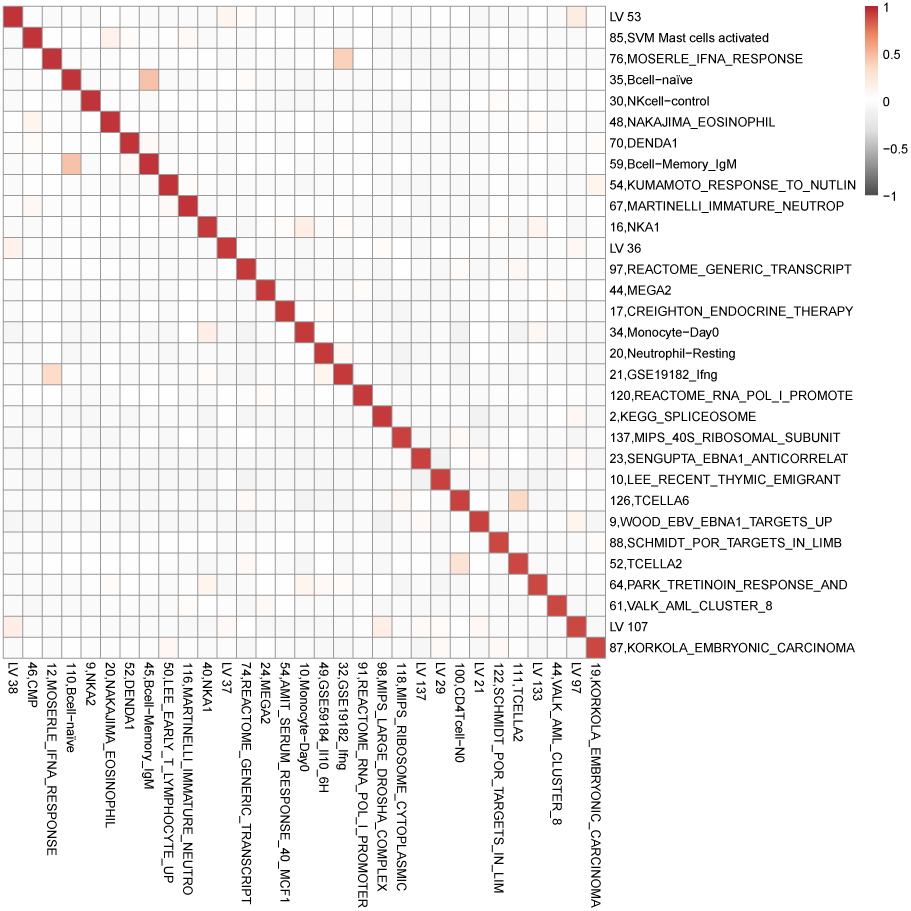
Heatmap of the correlations among LVs from decompositions performed on two different versions of the same data, one normalized for all technical variables and another normalized with quantile normalization. The heatmap shows all pairwise correlations for the top (rank correlation >0.9) best matched LVs named with their corresponding top prior information geneset(if any). Note that the prior information used is almost identical across the two decompositions.

Furthermore, when we compare the entire distribution of best matched correlations for LVs with- and without-prior information, as expected, LVs with prior information produce higher correlation matches supporting the hypothesis that these capture biological variation and are therefore relatively *normalization invariant* (Figure 7).

**Figure 7:**
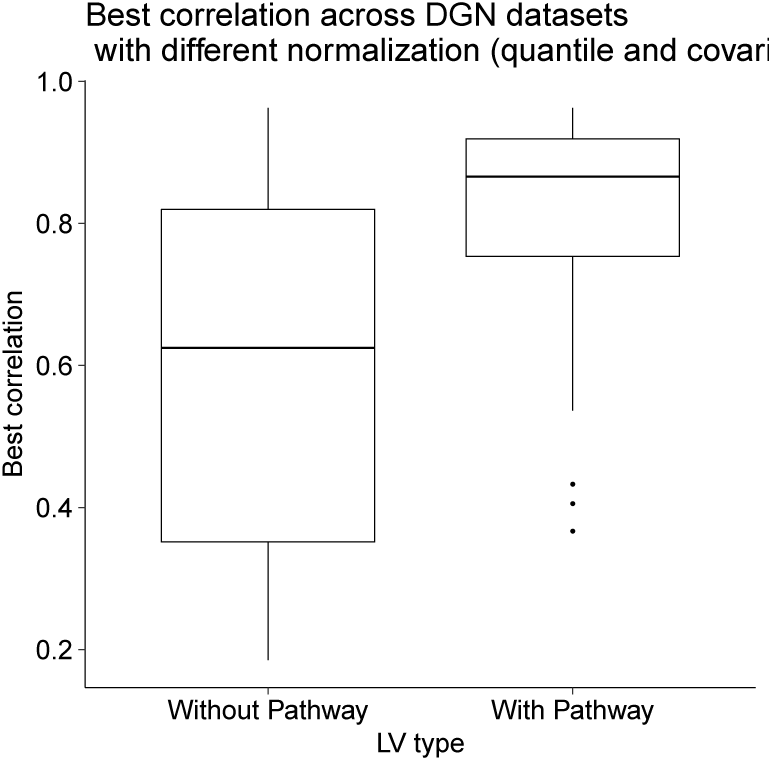
Correlation distributions across all best matched pairs of LVs estimated from two versions of the same dataset: one normalized with either all known technical factors and the other normalized by quantile normalization (see text). LVs that use prior information are more robust to normalization procedure as they more likely to have a high correlation match across differently normalized datasets.

### 2.6 The complete set of pathway-associated LVs found in the DGN dataset

The *U* matrix corresponding to the complete set of pathway associated LVs is visualized in Figure 8. For space reasons only the top pathway is used in the heatmap. The complete list of all pathways, and their corresponding statistics, can be found in the Supplementary File 1.

## 2.7 Details of genotype associated LVs

### 2.7.1 LV interpretation and naming

The top genes contributing to each genotype associated LV are depicted in Figure 9. In many cases the identity of the genes and the corresponding PLIER pathway utilization (see *U* matrix visualized in the main paper) point to a clear cell-type effect (LV44, LV133, LV56) or a canonical pathway (LV21, LV40, LV97, LV120). In these cases the LVs can be interpreted as estimating the specific cell-type proportion or pathway-level effect and are named accordingly.

**Figure 9:**
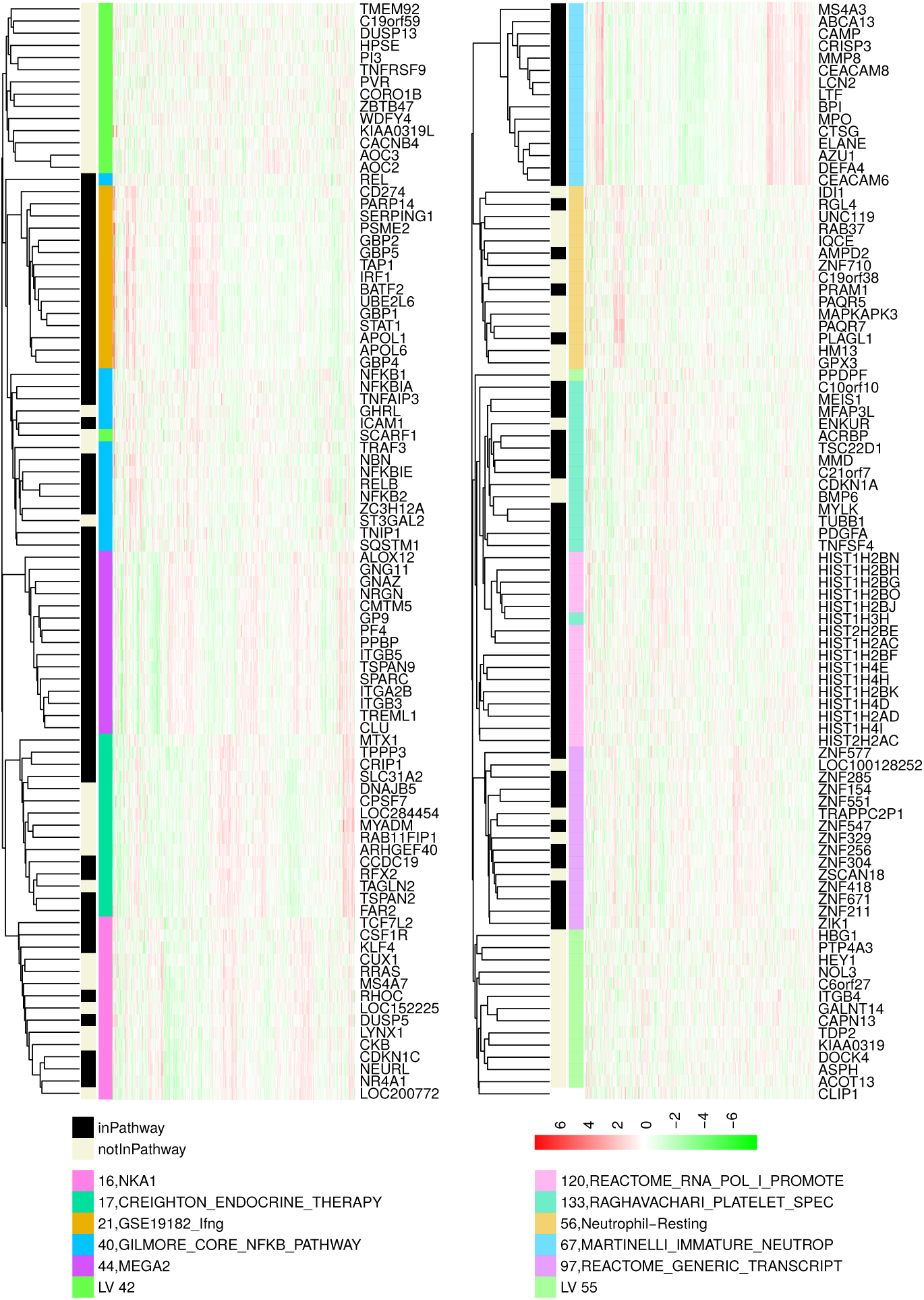
Top genes for all genotype associated variables.

In some cases the pathway utilization did not allow for unambiguous interpretations. For example, the top pathway for LV16 was “NKA1”, a NK-cell marker gene list. However the top genes in the LV loadings do not correspond to “canonical” NK-cell markers. This pattern is instead observed for LV30 which also makes use of NK pathways. Thus, LV16 cannot be interpreted as NK cell proportion though its pathway utilization suggests some relationship to NK cell biology. We also note that two of the LVs that had some of the strongest genotype associations did not use any pathway information. We hypothesize that collectively these LVs most likely represent transcription pathways that are not well annotated in our prior information though they may correlate with some prior information genesets.

Nevertheless, these transcriptional pathway potentially have some cell-type origin and we investigate this by checking the bias in cell-type expression in a large independent dataset of immune cell-types, ImmGen [Heng et al., 2008]. The results are visualized in Figure 10. We find that the top genes for LV16 were biased towards higher expression in myeloid and ILC cells which is consistent with being related to NK-type expression signature. LVs 17,42, and 56 were likewise biased towards myeloid cell-types. This is highly consistent with the the effects of the putative *cis* drivers (NEK6, PLAGL1, and IKZF1 respectively, see Table 1) on proportions of various myeloid cell-types as determined in a large GWAS study of blood cell-type composition [Astle et al., 2016]. LV55 had no identifiable signature in ImmGen data, however it is biased for genes expressed in the erythroid lineage based on DMAP (Differentiation Map) dataset [Novershtern et al., 2011]. Top genes include HGB1 (rank 5) and HGB2 (rank 16) – fetal hemoglobins that are expressed but not made into protein. Moreover the putative *cis* driver for LV55 eQTL is NFE2 which is a transcription factor known to be involved in erythrocyte and megakaryocyte development.

**Figure 10:**
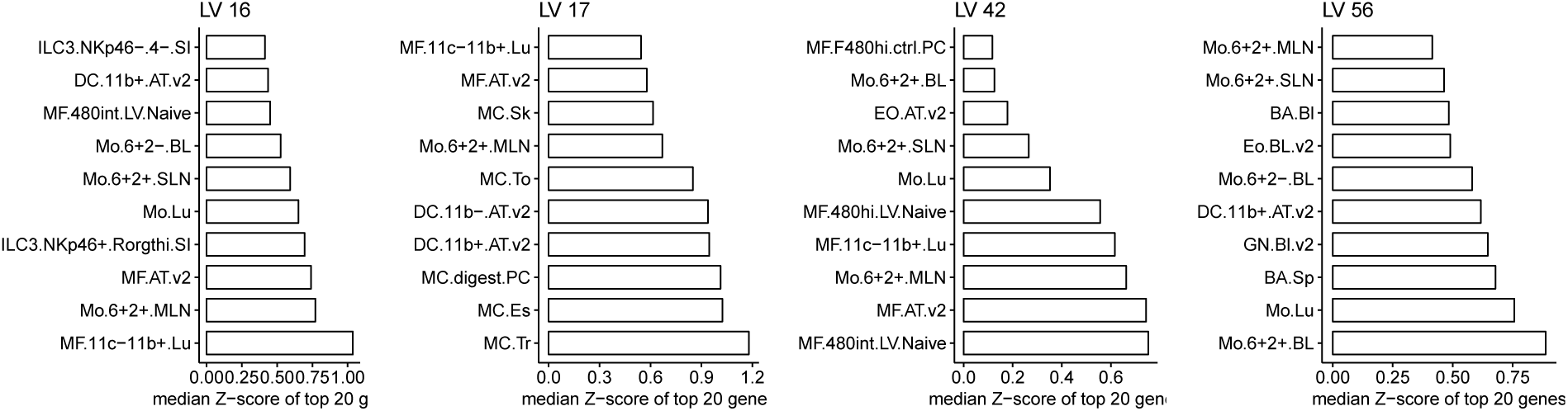
Top scoring Immgen cell-types for genotype associated LVs with no or ambiguous PLIER pathway annotation.

### 2.7.2 Independence of Megakaryocyte LVs

One of the key findings of this study is that LV44 and LV133 are both related to Megakaryocyte/platlet biology but are *independently* regulated by the ARHGEF3 locus. We find that while the estimates for these two LVs are correlated, the eQTL effects are substantially improved when regressing one LV on the other and using the residuals for eQTL testing (Figure 11).

## 3 Discussion

### 3.1 On the use of PLIER for mixture proportion estimation

We show that PLIER is capable of outperforming the best available model-based method (Cibersort) on mixture proportion estimation. Cibersort relies on known quantitative cell-type signatures, while SVM based framework is robust to outliers and discrepancies, it is likely that the hard-coded Cibersort signature is not a good fit for our dataset. Even though the cell-type marker genesets used by PLIER are in part produced from the same source data [Abbas et al., 2009, Novershtern et al., 2011], there are two important distinctions. PLIER is considerably more tolerant of errors in marker genes since the the model simply stipulates that we wish to find latent variables such that the loading on the marker genes are higher *on-average* than the background, without specifying a target value. Moreover, since PLIER automatically selects a few relevant pathways out of hundreds or thousands of available ones it can be supplied with multiple and possibly discordant markers sets for the same cell-type.

It is important to note that purpose of PLIER is general pathway-activity estimation. We do not expect that PLIER will substitute model-based methods for the explicit task of mixture component inference where model-based methods have several conceptual advantages. For example, PLIER operates best on z-scored data and thus by default discards valuable information about total transcript abundance. Moreover, PLIER is only applicable to relatively large datasets. In particular the number of major variance components, that can not be greater than the number of samples (and is typically much less), must be at least the number of mixture components we would like to estimate. Thus, PLIER cannot be applied for mixture component estimation in datasets with just a few samples, where model-based methods should have a clear advantage. Importantly, performance of model-based methods is highly dependent on the basis signatures (pure cell expression states) which may vary according to assay platform and processing pipe-line. A basis signature optimized for a particular data acquisition framework will provide optimal performance.

### 3.2 Alternative approaches

There are several methods that can take prior information about genesets into account in order to learn a biologically meaningful low dimensional representation. Examples, include Bayesian Factor Analysis [Bunte et al., 2016] that extracts pathway-level latent variables and our previously proposed method CellCODE [Chikina et al., 2015] that estimates cell proportion variation from cell-type marker genesets. However, these methods require that the genesets be specified *a priori* and that genes can be partitioned into these sets (though some overlap is allowed). In contrast, in our method the pathways themselves are subject to optimization and our method is designed to effectively choose just a few relevant genesets from thousands of available ones.

As our goal is to force gene loading to be represented by biologically coherent genesets it is natural to seek a solution based on group lasso regularization, which can perform variable selection at the group level. However, given that the biological genesets are highly redundant and overlapping, group lasso, which requires non-overlapping groups, is unsuitable. While it is possible to define more complex norms that accommodate group overlaps these have drawbacks. For example, a related method termed structured sparse PCA [Jenatton et al., 2009] has been developed for image analysis. This method implements a direct optimization of the column support, but can only constrain the support to be the complement of a union of predefined groups, which corresponds to rectangle-bounded regions for images, but is not interpretable for genesets. Another related method that considers biological genesets explicitly is the Overlap Group Lasso which employs an alternative norm that enforces the biologically desirable union-of-groups support [Obozinski et al., 2011]. However, the implementation is computationally expensive on large numbers of groups and in its native form does not explicitly deal the issue of geneset/pathway incompleteness.

### 3.3 Future developments

Despite the promising results there are a number of areas for potential improvement and future work will center on improving the recovery of LV with only a few supporting genes as well improving performance on very large geneset collections. For example, even on simulated data we find that increasing the amount of irrelevant prior information degrades the method’s performance. On the other hand, the available prior information represented in geneset databases such as mSigDB is constantly increasing which makes robustness to large prior information collections a top development priority.

## 4 Methods

### 4.1 Problem Setting

Given a gene expression profile *Y* ∈ ℝ^*n×p*^, where *n* is the number of genes and *p* is the number of samples, we state the original PCA as a matrix approximation problem. Suppose *n* > *k*, *p* > *k*. We wish to find *Z*, *B* minimizing

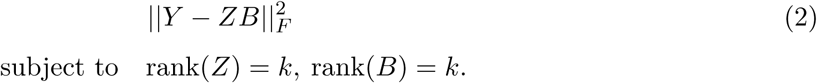

Since gene expression measurements are highly correlated, it is reasonable to expect that the data *Y* can be efficiently represented in this low dimensional space. Without imposing additional constraints on *Z* and B, an optimal solution can be obtained from the singular value decomposition (SVD) of *Y*. In an SVD based decomposition, rows of *B* are referred to as principle components (PCs). Since PCs are necessarily orthogonal, which our method do not require, we will use the more general term latent variables (LVs).

In order to improve the interpretability of the low dimensional representation in the context of known biology, we impose additional constraints on the matrix *Z*. Our aim is to encourage the loadings (columns of *Z*) to align as much as possible with existing prior knowledge. In the most general case such prior knowledge can be expressed as a series of genesets representing biological pathways, sets of tissue- or cell-type specific markers, and coordinated transcriptional responses observed in genome wide experiments.

Given *n* genes and *m* genesets, we represent the prior knowledge as a matrix *C* ∈ {0,1}^*n×m*^, so that *C_ij_* = 1 indicates that gene *i* is part of the *j_th_* geneset. Using the same notation as above, we define the revised decomposition problem based on the original formulation. We wish to find *U, Z, B* minimizing

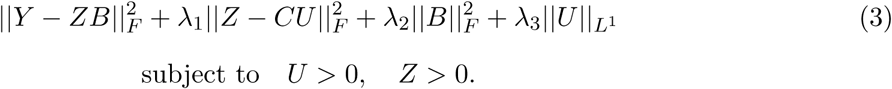

The first term of the optimization is the same as equation (1) and minimizes the overall reconstruction error. The second term specifies that *Z* should be “close to” sparse combinations of genesets represented by *C*. The third term introduces an *L^2^* penalty on B, while the fourth term is an L^1^ penalty on *U* (applied column-wise), which ensures that only a small number of genesets represent each LV.

The parameter *λ*_1_ keeps a balance between the proportion of prior knowledge we include and the degree to which we reconstruct the gene expression profile. We also restrict *U* and *Z* to be positive, which enforces that genes belonging to a single geneset are positively correlated with each other and the loadings are positively correlated with the prior information.

We solve the optimization problem by using block coordinate minimization, which iteratively minimizes the error on *Z*, *U*, and *B*. The complete method starts by initializing *Z* and *B* from the SVD decomposition and repeats the following steps until *B* converges.

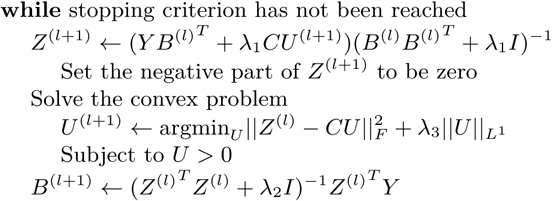

The stopping criterion is defined as a relative change in *B <* 5 × 10^−6^, or a leveling off in the decrease of the relative change in *B*. While there are no convergence guarantees, in practice this algorithm converges in under a few hundred iterations.

#### 4.1.1 Optimization constants

The optimization has 4 free parameters λ_1_, λ_2_, λ_3_, and *k* and internal cross validation cannot be used to optimize them as the reconstruction error 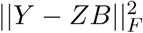 is always minimized when λ_1_ = 0. However, based on extensive testing with simulations and real data, we can set several default parameters that perform well in a range of situations. For example, we find that a reasonable starting value for *k* can be inferred from the the number of statistically significant PCs which can be determined via permutation by the approach proposed in [Leek et al., 2007] or the simple “elbow” approach (**num.pc** in our package implements both). However, it is logical that the number of constrained latent variables needed to explain the data is higher, and we suggest increasing the initial *k* by 50%. Importantly, the method is not sensitive to the exact value of *k*. LVs found at lower *k*’s persist when *k* is increased. It is also possible to keep increasing *k* as long as the number of LVs with prior information above some AUC and/or *p*-value threshold increases, but this requires multiple runs.

A good choice for λ_1_ and λ_2_ can be derived from the observation that if we consider the SVD decomposition of *Y* as *UDV^T^* we should have that *Z* ≈D^1/2^ and *B* ≈ D^1/2^V^T^. Therefore the diagonal elements of *Z^T^Z* and *BB^T^* are well approximated by *D* which thus gives the correct range for the relevant constants. By default we set λ_2_ = *d_k_* and λ_1_ = *d_k_/2* with the factor of 2 coming from the positivity thresholding on *Z*. It is also possible to optimize λ_1_ along with λ_2_ around its default value relative to some external validation source. For example, we can check how well the LVs recovered in *B* correlate with an independent dataset such as clinical variables, genotype, or another set of molecular measurements.

The correct value of constant λ_3_ that controls the sparsity of *U* is highly dataset dependent as it ultimately depends on how well the available prior information explains the data structure. We have devised an adaptive approach that works well for datasets of diverse characteristics. Specifically, we can specify the fraction of latent variables that we wish to be associated with prior information, 0.7 by default. The λ_3_ constant is then periodically adjusted by binary search to meet this goal. Even though this adaptive procedure keeps the number of positive entries in *U* constant regardless of prior information relevance, the significance of pathway association for each LV is ultimately tested by gene-holdout cross-validation (see bellow).

#### 4.1.2 Gene-holdout Cross Validation

It is natural to ask to what extent the non-zero coefficients of *U* represent non-random associations between loadings (columns of *Z*) and prior information. In order to quantify this we design a cross validation procedure that proceeds as follows. For each pathway included in the entire prior-information compendium a random 1/5th of the positive genes are set to 0 and this new prior information matrix is used to run PLIER. Afterwards, we can test how well the gene loadings in the PLIER output matrix *Z* are able to recover these held-out genes. Specifically, for each LV- pathway correspondence represented as a positive value in *U* we compute the AUC and p-value for the recovery of that pathway in the loadings of *Z* using the held-out set of genes as positive labels and genes not annotated to this pathway as negative labels. We find that the cross-validation procedure produces correct AUC estimates as *p*-values computed from a gene-level permuted prior information geneset ( which preserves dependencies among pathways) are uniformly distributed.

While this procedure necessarily discards some data and may adversely effect the ability to detect small pathways we find that the benefit of having accurate statistical estimates outweighs these concerns. PLIER will run in cross-validation mode by default but we allow for cross-validation to be turned off in which case all genes belonging to each geneset are used.

### 4.2 Validation Data

#### 4.2.1 Sample Processing

Blood was drawn into Tempus tubes (AB scientific) for RNA and into EDTA tubes for Cyto analysis respectively. RNA was extracted using the MagMAX^TM^ for Stabilized Blood Tubes RNA Isolation Kit (Fisher) following the manufacturer’s protocol. Libraries were constructed using the TruSeq Stranded mRNA kit (Illumina) at the Epigenetic core at the Weil Cornell medical college.

#### 4.2.2 CyTOF Sample Processing

CyTOF antibodies were either purchased pre-conjugated from Fluidigm (formerly DVS Sciences) or purchased purified and conjugated in-house using MaxPar X8 Polymer Kits (Fluidigm) according to the manufacturer’s instructions. Whole blood samples were processed within 4hrs of collection and stained by additional of a titrated panel of antibodies (table X) directly to 400uL of whole blood. After 20 minutes of incubation at room temperature, the samples were treated with 4mL of BD FACSLyse and incubated for a further 10mins. The samples were then washed and incubated in 0.125nM Ir intercalator (Fluidigm) diluted in PBS containing 2% formaldehyde, and stored at 4oC until acquisition.

Immediately prior to acquisition, samples were washed once with PBS, once with de-ionized water and then resuspended at a concentration of 1 million cells/ml in deionized water containing a 1/20 dilution of EQ 4 Element Beads (Fluidigm). The samples were acquired on a CyTOF2 (Fluidigm) at an event rate of ¡500events/second.

#### 4.2.3 CyTOF Data Analysis

After acquisition, the data were normalized using a bead-based normalization in the CyTOF software and uploaded to Cytobank for initial data processing. The data were gated to exclude residual normalization beads, debris, and doublets, and exported for subsequent clustering and high dimensional analyses.

Individual samples were first clustered using Phenograph [Levine et al., 2015], an agnostic clustering method that utilizes the graph-based Louvain algorithm for community detection and identifies a hierarchical structure of distinct phenotypic communities. The communities were then meta-clustered using Phenograph to group analogous populations across patients. These metaclustered populations were then manually annotated based on similar canonical marker expression patterns consistent with known immune cell populations. These annotations are also used to generate a consistent cluster hierarchy and structure across all samples in the dataset.

#### 4.2.4 RNAseq methods

RNA samples were sequenced SE100 to an average depth of 48.8 million reads. Quality assessment was done with FastQC [Bioinformatics, 2011]. Alignment to GenCode hg38 was done using STAR [Dobin et al., 2013]. Transcript counts are assigned using the FeatureCounts tool (subread package [Liao et al., 2013]).

### 4.3 Details of methods comparisons

We compare performance on the validation dataset against 4 alternative approaches, sparse principle component analysis (SPC), non-negative matrix factorization (MNF), seeded NMF, and Cibersort. For seeded NMF we first computed an undirected NMF decomposition and then replaced 30 of the *W* components with the LM22 Cibersort signatures (rescaled to match the range) and restarted the NMF optimization from this new *W* value. For NMF we used the default algorithm and matrix norm (Forbenius) and since it required a positive matrix we used the log2(FPKM) values which achieves better performance than raw FPKM values. SPC has no restrictions on the input and in our experiments performed best on z-scored data (z-scored data are also used for PLIER). The hyperparameters for SPC were set with cross-validation separately for each component as described in the original paper [Witten et al., 2009]. Since SPC and NMF do not assign a biological, causeto the inferred latent variables, for the purpose of evaluation the latent variables were matched to the single cell-type that they predicted best.

For Cibersort we used raw FPKM values, as suggested in the documentation, and the resulting estimates were matched to cell-type measurements according to cell-type identity. To account for the fact that our cell-type classes are slightly different from those predicted for Cibersort we allowed various combinations of Cibersort estimates, for example we created an “all-Bcell” estimate by adding naive and memory B cells and picked the best correlated estimate out of the three. A similar approach was taken for NK and dendritic cells for which Cibersort provides separate activated estimates. We also created several combinations of Cibersort CD4 estimated to represent a possible estimate of our activated (CD45RA-) measurement.

### 4.4 Public Data

#### 4.4.1 DGN dataset

The Depression Gene Networks (DGN) dataset is not available for public release but can be requested from National Institute of Mental Health (NIMH) following instructions in the original publication [Mostafavi et al., 2014]. The NIMH database contains several normalized versions of this data and for our study we used “trans” normalized data as described in [Battle et al., 2014]. This data is already normalized for genotype principle components and all known technical factors and no further normalizations were performed.

#### 4.4.2 NESDA dataset

The NESDA dataset was normalized for batch effects using th HCP method described in [Battle et al., 2014]. The resulting data was further normalized for the first 5 genotype principle components using linear regression.

#### 4.4.3 Prior information genesets

The generic blood cell-type marker dataset was derived from the IRIS (Immune Response In Silico) [Abbas et al., 2009] and DMAP (Differentiation Map) datasets [Novershtern et al., 2011] datasets. Many canonical marker genes (such as CD19, CD3E, CD8A) have a multimodal distribution with on high expressor group and one or more low/medium expressor ones. The highest expression group typically does not overlap with lower expression distributions and we base our marker selection metric on this observation. Genes were considered to be markers if they could be partitioned into high and medium/low expression so that the difference between minimum and maximum values respectively (the gap between these distributions) exceeds a threshold (we used 2 for IRIS and 0.7 for DMAP). This procedure results in highly overlapping sets of markers for related cell-types however our method is flexible and can easily handle redundancy. The marker sets derived from the IRIS and DMAP datasets are included in the PLIER R package. For the purpose of analyzing DGN we also included cell-type markers from a recent publication [Newman et al., 2015] which covers fewer cell-types but with highly optimized marker sets. The complete prior information dataset used for DGN analysis includes cell-type markers, “canonical pathways” and “chemical and genetic perturbation” genesets from mSigDB, and a set of transcriptional signatures relevant to immune signaling described in [Filiano et al., 2016].

### 4.5 trans-eQTLs

Trans-eQTLs effects in the DGN dataset were assessed with Spearman rank correlations and were Bonferroni adjusted by the total number of tested SNPs (651075). To assess replication in the NESDA dataset SNPs were matched based on LD using the LDlink tool with CEU population [Machiela and Chanock, 2015]. Specifically, if the exact SNP was not present in the NESDA dataset we selected the SNP with the highest LD, and if multiple SNPs had the same LD, we took the one closes in genomic coordinates. We only considered a match if the best LD was above 0.8. To asses *π_1_* in a consistent manner we used a single threshold of Λ = 0.5. Using a single threshold the *π_1_* is computed as the fraction of *p*-values bellow 0.5 divided by 0.5.

### 4.6 Platelet phenotypes

The sentinel SNPs and their relevant phenotypes (MPV, PLT, or both) are supplied as the supplementary table in [Gieger et al., 2011]. Proxy SNPs were defined as above. SNPs were deemed significant if they had a Bonferroni corrected p-value < 0.05 where the Bonferroni factor is 80 (total number of SNPs with platlet phenotypes tested).

